# Delayed normalization model captures disparate nonlinear neural dynamics measured with different techniques in macaque and human V1

**DOI:** 10.1101/2023.01.30.525700

**Authors:** Jingyang Zhou, Matt Whitmire, Yuzhi Chen, Eyal Seidemann

## Abstract

Neuronal representations throughout the primate visual system display a wide range of nonlinear dynamics in response to static visual stimuli. To understand the neural basis of visual perception, it is important to quantify these nonlinearities and to develop general models that allow one to predict response dynamics to visual stimuli with arbitrary temporal waveform.

Voltage-sensitive dye imaging (VSDI) is a powerful method for measuring neural population responses from the cortex of awake, behaving, subjects. Here we used VSDI to measure the dynamics of neural population responses in macaque V1 to visual stimuli with a wide range of time courses.

We found that beyond clear nonlinearities for briefly presented visual stimuli, stimulus-evoked VSDI responses are surprisingly near additive in time. These results are qualitatively different from neural dynamics to similar stimuli previously measured in human visual cortex using fMRI and electrocorticography (ECoG), which show strong sub-additivity in time.

To test whether this discrepancy is specific to VSDI, a signal dominated by subthreshold neural activity, we repeated our measurements using a genetically encoded calcium indicator (GCaMP), a signal dominated by spiking activity. We found that GCaMP signals in macaque V1 are also near-additive. Therefore, the discrepancies in the degree of additivity between these different measurements are not attributed to the difference between sub- and supra-threshold neural response.

Finally, we show that a simple yet flexible delayed normalization model can capture the dynamics of all of these measurements, suggesting that dynamic gain-control is an important mechanism contributing to neural processing in the brain.

## Introduction

Neural representations throughout the primate visual system display a wide range of nonlinear dynamics to visual stimuli [1-7]. To understand the neuronal basis of visual perception, it is important to quantify the dynamics of neuronal responses and to characterize nonlinearities within these dynamics. This is because nonlinearities are central for our rich and flexible visual perception [8].

Neural representations in primate primary visual cortex (V1) are widely distributed: a small and localized visual stimulus activates millions of neurons distributed over more than 10 mm^2^ in a macaque’s V1 [9, 10]. No current technique can measure activities of all neurons that may be perceptually relevant with single-cell resolution and real-time dynamics in V1 of a behaving primate. Therefore, to understand the neural basis of visual perception, we need to combine multiple complementary techniques, each of which samples a different aspect of brain activity at a different spatial and temporal resolution. Here, we take a step towards this long-term goal, by comparing and contrasting neuronal dynamic responses to visual stimuli in primate visual cortex obtained with different measurement methods and across different spatial and temporal scales.

Neuronal dynamics measured using different methods can exhibit different properties. This is because different measurement methods emphasize different signals within a population of neurons. For example, widefield VSDI signals are dominated by spatially pooled membrane potentials of cortical neurons [9, 10], and widefield GCaMP signals appears to be more closely linked to pooled spiking activities [11]. fMRI is the most prevalent method used to measure behaving humans’ brain responses. fMRI is an indirect measure of neuronal activities, and it has been challenging to relate the fMRI BOLD signal to a local neuronal population’s stimulus-evoked responses [12-15].

In the current study we set out to use VSDI to measure population responses from V1 of behaving macaques to large and high-contrast visual stimuli that were flushed with a range of time courses. Our first goal was to assess the degree of nonlinearity in these responses and to develop a general computational model that can predict the temporal dynamics of V1 responses to a visual stimulus with any arbitrary temporal waveform.

Our second goal was to compare the nonlinearity in our VSDI results with the temporal additivity of fMRI and ECoG responses in human visual cortex [15, 16]. We found that stimulus-evoked dynamics measured using these methods differ qualitatively in their properties. Stimulus-evoked VSDI dynamics measured in behaving Macaques’ V1 are near-additive over a wide range of stimulus time courses while fMRI and ECoG dynamics measured for similar stimuli in human V1 show strong sub-additivity in time.

To test whether this difference is due to disparate spiking versus subthreshold neural population dynamics, we measured widefield GCaMP responses to identical stimulus conditions in behaving macaque V1. We found that GCaMP signals in macaque V1 are also near-additive, suggesting that the difference in sub-vs. supra-threshold dynamics are unlikely to be the source of the difference between the nonlinearities measured in fMRI, ECoG and in VSDI.

Finally, we show that a simple delayed normalization model can qualitatively account for the dynamics of all of these measurements, suggesting that dynamic gain-control is likely to be an important mechanism contributing to neural processing in the brain.

## Results

### Testing temporal additivity in stimulus-evoked VSDI dynamics

To assess properties of stimulus-evoked VSDI responses, we presented a set of large static patterned images (band-pass filtered noise peaking at 3 cpd, 100% contrast) with different temporal conditions (Figure 1A) while monkeys performed a fixation task. The first type of temporal condition is a single pulse of image presented using 6 different durations. The presentation durations range from 20 to 640 milliseconds, and the duration in each condition is twice as long as the previous condition (20, 40, 80, 160, 320, 640 ms). The second type of temporal condition is a single image presented twice, each for 160 ms with different inter-stimulus intervals (ISIs). The ISIs range from 20 to 640 milliseconds, and like varying durations, each ISI is twice as long as the previous ISI (Figure 1B). Because the stimulus is larger than the portion of the visual field represented in our imaging cranial windows, it elicits a response that is spatially uniform within the imaged area. We focus on analyzing the time course of the response within an area of several square mm within the center of the imaging chamber.

**Figure 1.**
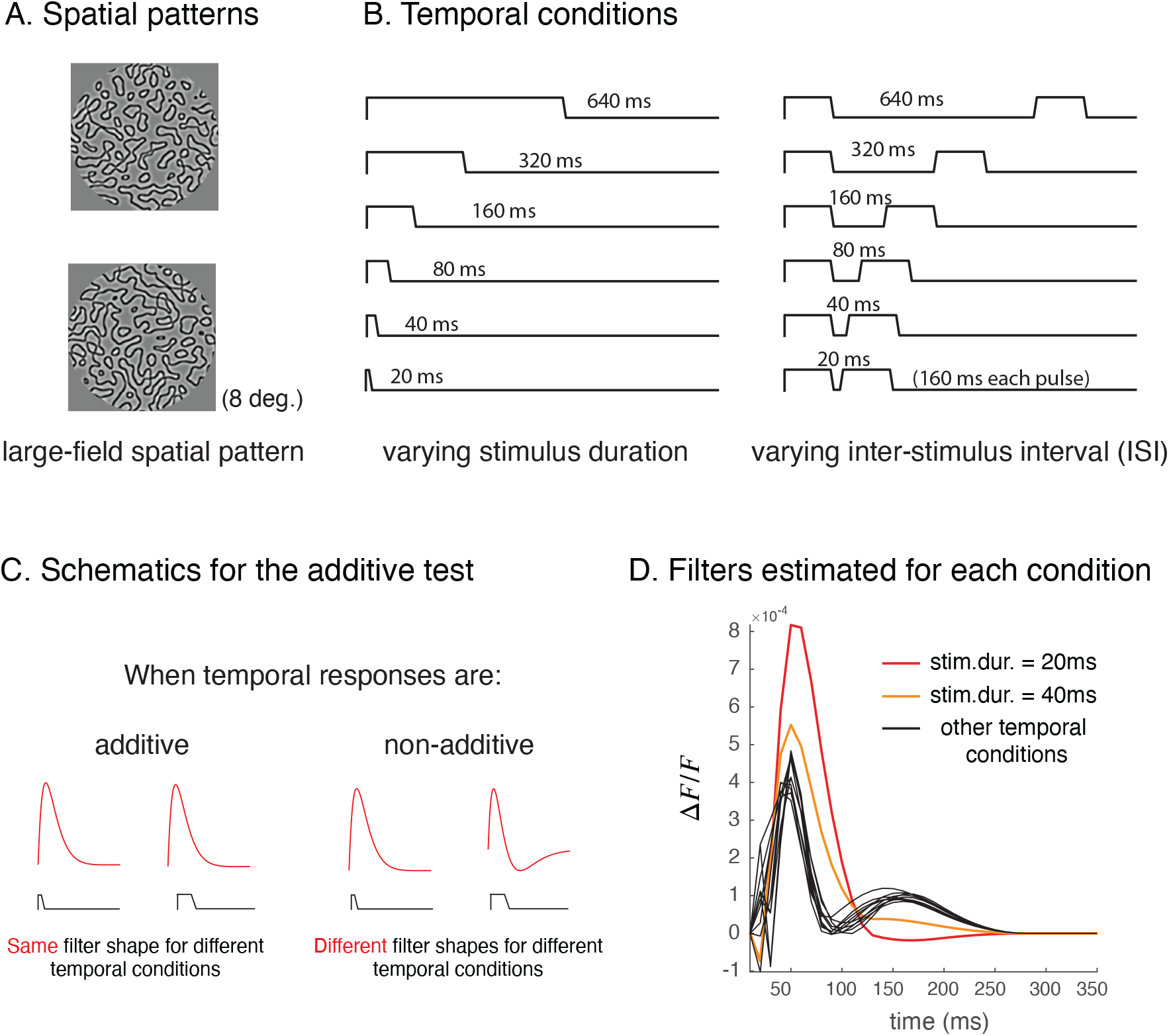
Stimulus and additivity test. A. Monkeys viewed large patterned stimuli that strongly activated population of V1 neurons. B. Subjects were presented with a single pulse of stimulus with 6 different durations, and double pulses of a single image with 6 different inter-stimulus-intervals (ISIs). C. If neuronal responses are linear (or are additive in our case), impulse response functions estimated from different temporal conditions would share the same shape. Stimulus-evoked responses are not linear if filters estimated for different conditions have different shapes. D. Besides when stimulus durations were brief, filters estimated from stimulus-evoked VSDI responses are relatively similar in shape. Each filter was estimated from the trial-averaged time course for a single stimulus condition.

To analyze stimulus-evoked VSDI dynamics, we first assess whether these dynamical responses to the two sets of temporal conditions can be explained by a linear model. Raw VSDI dynamics consist of two components [17, 18]. The first component is relatively fast, and closely tracks the stimulus time course *s*(*t*). This component is related to stimulus-evoked population membrane potential responses [19, 20]. We modeled this component using a linear filter *f*(*t*) convolved with the stimulus time course *s*(*t*). The second component of the VSDI time courses is a slow-varying compound signal that is likely to reflect a mixture of neural and non-neural slow variability. We modeled this component using a slow-varying function *g*(*t*), which is at least an order of magnitude slower than the first component [21, 22]. Overall, the linear prediction of the VSDI dynamics *r*_*l*_(*t*) can be summarized by additively combining the two components:

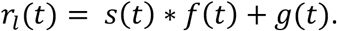

In Supplement, we show that *g*(*t*) is generally independent of stimulus conditions (Figure S1). Therefore, to understand how the fast stimulus-evoked component varies with the stimulus conditions, we removed *g*(*t*) from raw VSDI time courses, and used the remaining data for further analyses. For details of data processing, see Methods.

To quantify stimulus-evoked VSDI dynamics (the fast component), we first assess how and to what extent the extracted stimulus-evoked responses deviate from the predictions of a linear model. To do so, we fitted the linear model to trial-averaged data for each condition, and estimated a separate temporal filter for the stimulus-evoked component in response to each stimulus time course. If stimulus-evoked dynamics are linear, filters fit to different temporal conditions should share the same shape – a linear system can be uniquely characterized by a single filter (Figure 1C). If filters fit to different conditions have different shapes, this will indicate that the system is nonlinear and we can further study the types of nonlinearities that exist in the data (e.g. [23]) through observing these differences.

The VSDI filter shapes estimated for different stimulus conditions are similar to those estimated in single-cell membrane potentials [24-26]. Surprisingly, the estimated filters for all temporal conditions beyond the two shortest durations share the same shape, suggesting that the pooled membrane potential in V1 is nearly linear in time for stimulus duration equal to or longer than 80 ms. For brief stimulus durations (< 80 ms), estimated filters tend to be larger in gain (higher in amplitude), be monophasic rather than biphasic, and have slower dynamics (Figure 1D). For example, the estimated time to peak for filters were around 50 ms for brief stimulus durations, and were around 40 ms for longer-duration stimuli. Our observation confirmed that stimulus-evoked VSDI responses deviate from the linear prediction, and the deviation is possibly due to gain adjustment that depends on stimulus durations. This observation is consistent with duration-dependent gain adjustments observed in neuronal responses measured using other methods and in other parts of visual pathways [16, 27, 28]. In a later section, we will compare across measurement methods, how much temporal dynamics deviate from the linear predictions.

### Capturing non-linear VSD dynamics using a delayed normalization model

To account for gain changes in the stimulus-evoked VSDI responses to brief stimuli, we devised a delayed normalization model [4]. The model has a divisive form, and consists of a numerator and a denominator, each involves a linear computation. The model numerator consists of a filter *f*_1_(*t*) convolved with a stimulus time course, *s*(*t*) ⋇ *f*_1_(*t*). The model denominator consists of another filter *f*_2_(*t*) convolved with the same stimulus time course, *s*(*t*) ⋇ *f*_2_(*t*). A positive constant σ is added to the denominator to prevent computations from becoming undefined when stimulus time course is 0, which is the case when stimulus contrast is zero. The delayed normalization model intends to summarize the stimulus-evoked component in the VSDI signal. Moreover, we additively combined the delayed normalization with *g*(*t*) to account for slow variations in the measurement time series. We fit the combined model to the overall VSDI dynamics to reduce estimation bias, and the combined model prediction *r*_*n*_(*t*) has the following form:

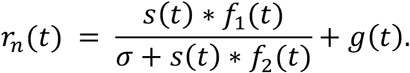

A single set of delayed normalization parameters {*f*_1_(*t*), *f*_2_(*t*), *σ*} were fit to all temporal conditions, and different sets of slow dynamics *g*(*t*) were fit to different temporal conditions in the trial-averaged data. This is because in the fast stimulus-evoked component, the same population of neurons are activated by different stimulus conditions. But the slow components, on the other hand is generally stimulus independent (see Methods and Figure S2).

To understand whether and to what extent the delayed normalization model improved upon a linear model, we compared the delayed normalization model to the linear model with a single filter fit across all temporal conditions (Figure 2A). The delayed normalization model improved upon the linear model, especially when stimulus durations are brief (Figure 2B). We can infer why the delayed normalization model improves the fit by examining how the model works. At stimulus onsets, the model numerator starts to respond, and this linear computation dominates the response dynamics before normalization dynamics kick in. Because the denominator filter is delayed compared to the numerator filter, after a brief time period, the normalization dynamics start to strengthen, and decrease the response gain while speeding up the dynamics. The temporal difference between the stimulus-drive (numerator) and the normalization drive (denominator) can account for our stimulus-evoked VSDI responses – dynamics are slower and the gain is higher at brief stimulus presentations.

**Figure 2.**
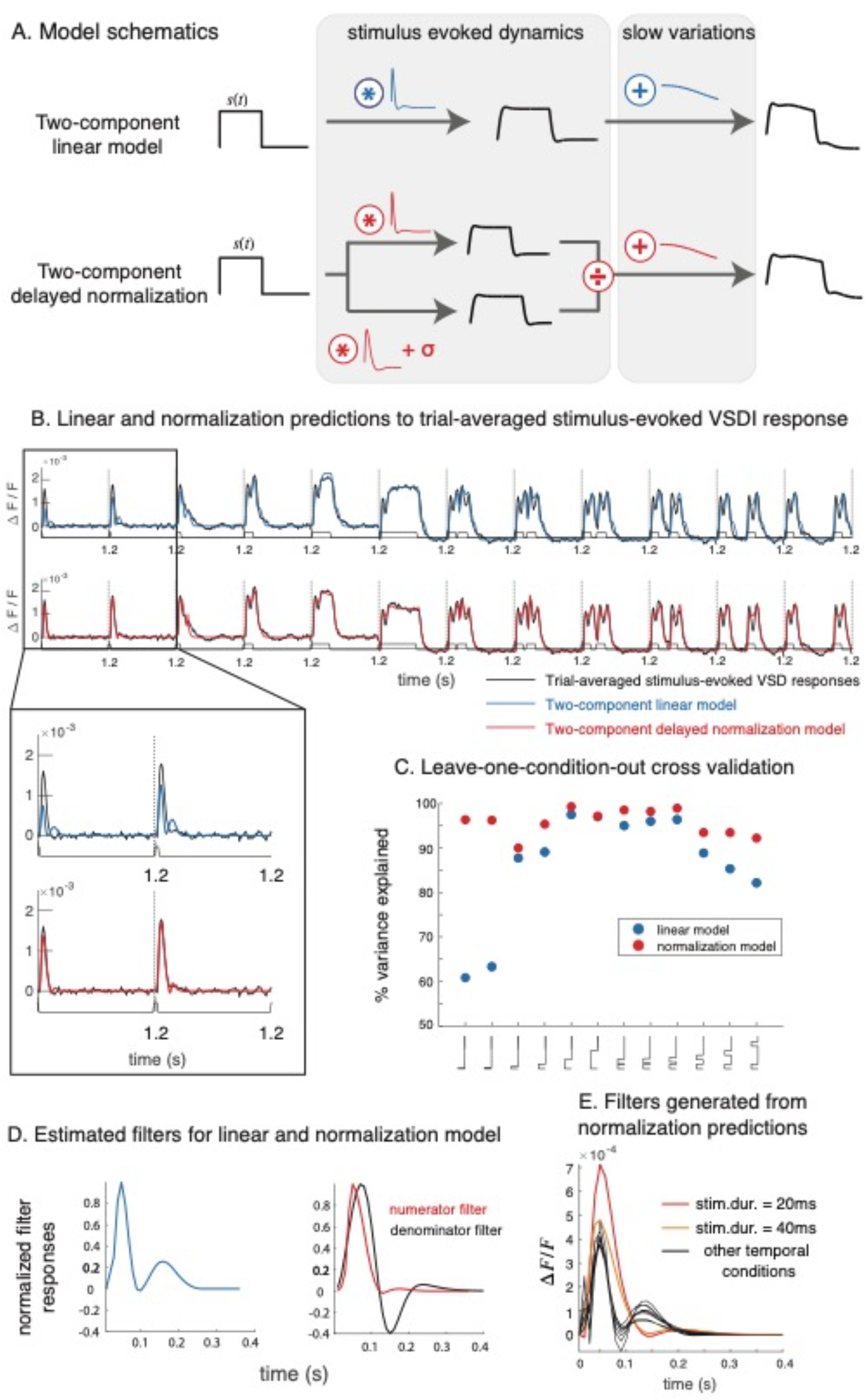
Fitting linear and delayed normalization models to VSDI dynamics. A. The linear model consists of a fast component (a filter convolved with a stimulus time course) that captures stimulus-evoked dynamics. The delayed normalization also consists of a divisive computation that intends to capture the same dynamics in VSDI measurements. B. Delayed normalization model, compared to the linear model, better captures stimulus-evoked VSDI dynamics when stimulus presentations are brief. C. Leave-one-condition-out cross validation further confirms that the delayed normalization improved upon the linear model at brief stimulus durations. D. Estimated filter for linear, and for normalization model (numerator filter). Both the linear filter (blue), and the numerator filter (in stimulus-drive) for the normalization model (red) are relatively similar in terms of time to peak, but the numerator filter in the normalization model tends to be simpler in shape. This is because some of the dynamics captured by the linear filter was absorbed into the normalization dynamics. The filter time to peak for the normalization dynamics (denominator of the delayed normalization model) is delayed compared to the numerator filter, and the denominator filter is biphasic in shape (black). E. Linear filters estimated from the normalization predictions to the stimulus-evoked VSDI component for each stimulus condition closely resemble what we observed in Figure 1.

The delayed normalization model performed better than the linear model in leave-one-condition-out cross-validations (Figure 2C). This type of cross-validation examined whether fitted model parameters well-generalize to predict response dynamics in a different stimulus condition. We further investigated the filter shape estimated using delayed normalization (Figure 2D). The filter estimated for the stimulus-drive (numerator filter) is comparable to the linear model filter in terms of the time to peak (50 ms for both the linear model filter and the delayed normalization numerator filter). Compared to the linear model filter, the numerator filter of the normalization model tends to have a simpler (and more monophasic) shape. This is because some dynamics of the linear model filter were absorbed into the normalization dynamics. In the delayed normalization model, the peak of the denominator filter is around 70 ms, which is delayed compared to the numerator filter time-to-peak, and the estimated denominator filter has a biphasic shape. Using the predictions of the normalization model, we estimated one linear filter for each temporal condition as we’ve done in Figure 1 (Figure 2E). The estimated filter shapes for all stimulus conditions closely resemble what we estimated directly from the VSDI data.

### Comparing temporal additivity measured using other methods

In the previous section, we accounted for stimulus-evoked VSDI dynamics using a delayed normalization model. Delayed normalization has also been applied to describe dynamics measured using fMRI and ECoG responses in human subjects’ visual cortices [16]. Surprisingly, we found that stimulus-evoked VSDI dynamics qualitatively differ from those measured using these other two methods. Stimulus-evoked VSDI dynamics are near-additive in time beyond brief stimulus presentations, whereas dynamics measured using fMRI and ECoG (broadband signals that exhibit properties consistent with fMRI measures) are substantially sub-additive in time in V1 across all tested stimulus durations.

To compare additivity across measurement modalities, we adopted a different metric of additivity that can be applied to measurements with slow sampling rates. For example, it becomes extremely challenging to measure neuronal filter shapes using fMRI [15, 29], because the BOLD signals are sluggish and are typically sampled every 1 – 2 seconds. To compute this new additivity metric, we integrated the time course of the stimulus-evoked VSDI component within each condition to obtain twelve numbers, one for each condition (Figure 3A, also see Methods). If stimulus-evoked dynamics are additive, doubling stimulus duration doubles the sum of the response dynamics, and varying inter-stimulus intervals would not change the summed response [15]. Stimulus-evoked VSDI signals closely follow the additive prediction, consistent with the nearly identical shapes of the linear filters for stimulus duration at or above 80 ms (Fig. 1D). This near-additive response is further confirmed in a second data set, see Supplementary Figure S3.

**Figure 3.**
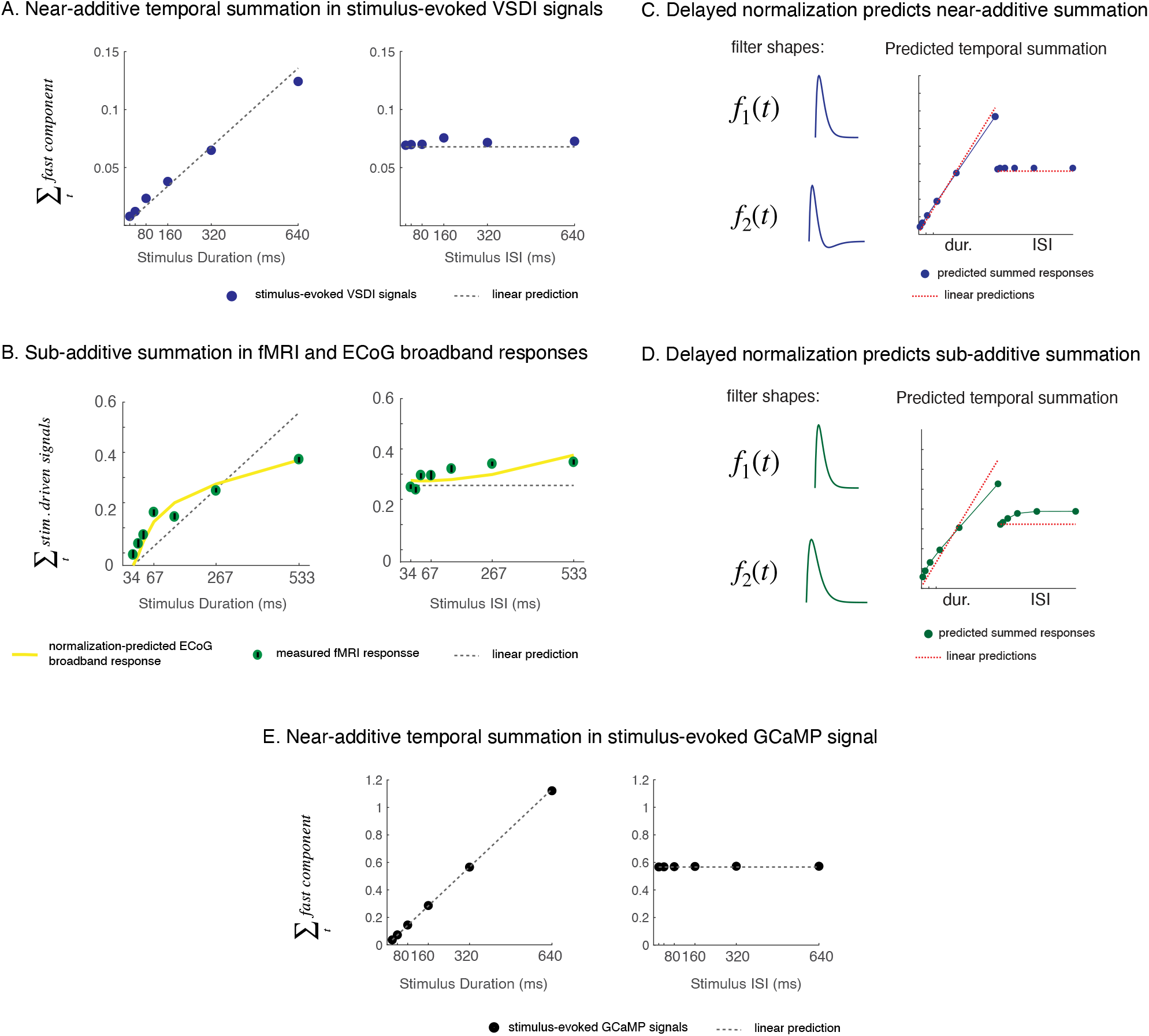
We used a different additivity test (as in [15]) that allows us to compare temporal additivity across measurement modalities. A. In VSDI, for example, we summed up the stimulus-evoked signals within each condition. Each black dot represents the response sum for a temporal condition, and the dotted line represents the linear prediction. Stimulus-evoked VSDI signals are near-additive. B. fMRI (yellow) and ECoG broadband signals, on the other hand, are substantially sub-additive: for example, doubling stimulus duration increases the summed response, but the increase is less than doubled (adopted from Ref [15]). C, D. The normalization filters estimated for ECoG data are monophasic (low-pass), whereas the normalization filters for VSDI data tend to be biphasic (band-pass) [16]. Using monophasic versus biphasic normalization filters, we can generate sub-additive and near-additive temporal dynamics. E. Stimulus-evoked widefield GCaMP responses from behaving macaque V1 (black dots), like VSDI signals, are also near-additive.

fMRI signals are thought to reflect activities of a local population of neurons, and have been demonstrated to be highly correlated with both local population spike rates and local field potentials (LFP) – slow electrical signals that include, but are not limited to, synaptic potentials [12, 14, 30, 31]. Even though fMRI measurement is coarse in temporal resolution, it can still capture the total response sum (but not dynamics) relating to neuronal activities evoked by a stimulus presented over different time courses. With fMRI data, a previous study [15] tested temporal additivity using a similar set of varying duration and ISI conditions (same type of stimulus patterns, response averaged across 4 human subjects). If the underlying neuronal signals vary additively in time, doubling stimulus duration would double the amplitude of fMRI response. Cortical responses measured using fMRI are substantially sub-additive. In particular, response amplitude increases but less than doubles when stimulus duration doubles. In the two-pulse temporal conditions, amplitude of the fMRI response to the second stimulus pulse also depends on ISI – response to the second pulse is suppressed for short ISIs and recovers over longer ISIs (Figure 3B). An ECoG data set (broadband signals, 70-210 Hz range, chosen because this range correlates with local multi-unit activities near the electrodes) is consistent with the fMRI temporal sub-additivity [16] (Figure 3B).

Delayed normalization model paired with different filter shapes can account for both near-additive and sub-additive temporal summation. In VSDI, the measured normalization filter (in the denominator) was biphasic, whereas the normalization filter used to account for ECoG data, which were substantially sub-additive as fMRI BOLD signals, was monophasic [16, 32]. In Figure 3C and 3D, we demonstrated that with different normalization filter shapes, the model could account for both near-additive and sub-additive temporal summations. Monophasic normalization filter suppresses the input drive over the entire summation period, whereas the suppressive effect ends at the end of the summation period for a biphasic filter.

One possible source for the qualitatively different temporal additivity of VSDI and fMRI signals could be the different nature of the neural signals that these methods are related to. VSDI captures the dynamics of population membrane potentials (mostly sub-threshold), whereas fMRI BOLD signals may be correlated mostly with spiking activities (supra-thresholds) (e.g. [33]). To examine this possibility, we ran the same temporal experiment using an additional measurement modality -- widefield imaging of genetically encoded calcium indicator (GCaMP6f) in V1 of a behaving macaque [11]. Widefield GCaMP signals sample neural signals at spatial and temporal scales similar to widefield VSDI, and were shown to be approximately linearly relate to summed local spiking activity [11]. If the differences between VSDI and fMRI temporal additivities are due to differences between membrane potential and spiking dynamics, we would expect GcaMP signals to be sub-additive as the fMRI signals. However, we found that GCaMP signals are like VSDI, and are even slightly more additive compared to stimulus-evoked VSDI signals (Figure 3C). For GCaMP filter shapes, and additional comparison between VSDI and GCaMP signals, see supplementary Figure S4. Therefore, our GCaMP results suggest that temporal nonlinearities of sub- and supra-threshold population responses in V1 are similar, and that other factors must account for the qualitative difference between VSDI and fMRI dynamics.

## Discussion

Here, we described stimulus-evoked VSDI dynamics using a generalized delayed normalization model. Compared to the linear model, delayed normalization accounts for both higher gain, and slower dynamics in the data when stimulus presentations are brief. Additionally, we compared stimulus-evoked dynamics across different population measurement methods. Surprisingly, we found that stimulus-evoked VSDI and GCaMP dynamics in macaque V1 are near-additive, but fMRI signals measured using comparable stimulus conditions in human V1 are substantially more sub-additive.

In general, deviations from linearity can be partitioned into two types. The deviations can either come from a lack of additivity – doubling the stimulus duration does not result in twice the total amount of responses. Or the deviations can be a result of lacking homogeneity – doubling the stimulus amplitude (i.e. stimulus contrast in our case) does not double the total response. We used delayed normalization to account for deviations from additivity in stimulus-evoked VSDI signals. In previous works, VSDI dynamics were also observed to deviate from homogeneity [34]. Here, we demonstrate that the delayed normalization model, with its parameters fit to VSDI response to different temporal conditions, can also qualitatively capture VSDI deviations from homogeneity.

Sit et al. 2009 [34] observed that when doubling the contrast of a stimulus time course, the stimulus-evoked VSDI responses increase but are less-than-doubled, and the dynamics of the signal also become faster. To demonstrate that the delayed normalization can encompass these previous observations, we used the model parameters fit to trial-averaged VSDI time courses to generate predictions to a single pulse of stimulus time course (200 ms) presented with different contrasts (Figure 4A). We generated five different contrast levels, by multiplying the stimulus time course with 5 different scalars, each indicating a contrast ranging from 6.25% to 100%. Predictions of delayed normalization qualitatively agree with previous observations: the predicted response gains are different at different contrast (logarithmic increase in contrast results in a near-linear increase in response amplitude) (Figure 4B), and shapes of the response dynamics are also different at different contrast levels (Figure 4C).

**Figure 4.**
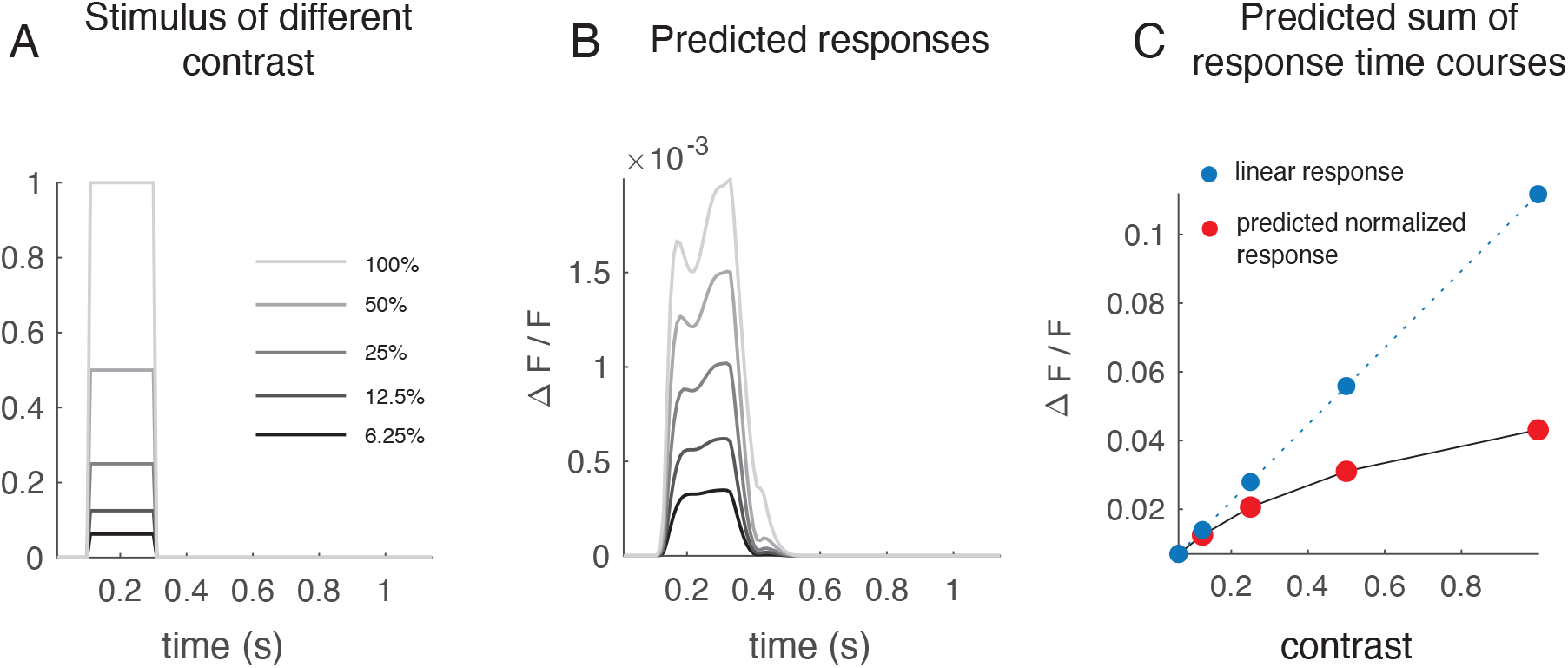
Delayed normalization can qualitatively account for deviation from homogeneity measured in stimulus-evoked VSD signals. A. We simulated stimulus-evoked VSD response to a 400ms stimulus presentation scaled with different contrasts, ranging from 6.25% to 100%. We used parameters fit to trial-averaged responses to different temporal conditions for this simulation. B. The simulated response time courses for different contrast differ in shape. C. Based on this simulation, doubling stimulus contrast responses in less than doubled total responses, consistent with previous observations.

In Results, we compared additivity across measurement methods, and found that properties of temporal dynamics measured in stimulus-evoked VSDI and GCaMP signals qualitatively differ from that in fMRI signals. In particular, the stimulus-evoked VSDI and GCaMP signals are near-additive except during a very brief stimulus presentation interval (< 80ms), whereas fMRI and ECoG broadband signals were sub-additive over a time range of hundreds of milliseconds. This conclusion is robust, and depends little on how we extracted stimulus-evoked dynamics in VSD and GCaMP measurements. To illustrate the robustness of our conclusion, we devised additivity tests using other extraction methods from the literature (Figure S5). Existing methods generally assume raw VSDI (and GCaMP) time courses consist of two additively combined components -- a fast component that reflects stimulus-evoked responses, and a slow “trend” component [18]. Our extraction method, compared to existing methods, assumes a more flexible form for the slow-component, and a delayed normalization model for the fast component (see Methods). Stimulus-evoked signals in VSDI and GCaMP extracted using these other methods are slightly noisier, but we can reach the conclusion of near-additive temporal summation (Figure S5). For details and implementation of each extraction model, see Methods section.

Delayed normalization was fit to stimulus-evoked VSDI components in our current paper, and was fit to ECoG broadband (70 Hz – 210 Hz) dynamics in a previous analysis. First of all, ECoG broadband signals were thought to better correlate with local population spiking responses. ECoG broadband excludes signals in the low-frequency range (<70 Hz), whereas VSDI measurements are dominated by the low-frequency signals. This difference in sampling frequency range can potentially provide one explanation to the difference of dynamics between the two signal types. ECoG broadband, once combined with the lower-range frequencies, could potentially produce signals that are near-additive in time. Second, ECoG broadband signals and VSDI signals have very different dynamics: for a sustained stimulus time course, ECoG broadband signals respond with a transient followed by a decay [16], whereas VSDI dynamics do not decay for prolonged stimulus presentations. This dynamical difference can potentially contribute to the near-additive versus sub-additive temporal properties in the two signal types. Examining filters fit to both data types, we found that for ECoG broadband signals, both the numerator and the denominator filter in delayed normalization tend to be monophasic. But for VSDI dynamics, the estimated denominator filter tends to be biphasic. In other words, the difference in temporal properties of the two measurements can be accounted by different normalization dynamics. Whether this difference in estimated filter shapes has any mechanistic grounding awaits to be further examined, but with a delayed normalization model, we can quantify the difference, and generate predictions to new stimulus conditions based on these parametric differences.

## Methods

### Data collection

All procedures have been approved by the University of Texas Institutional Animal Care and Use Committee and conform to NIH standards. The experimental techniques for optical imaging in behaving monkeys were similar to what was described in [10]. In brief, a metal head post was implanted for each animal, and a metal recording chamber located over the dorsal portion of V1, a region representing the lower contralateral visual field at eccentricities of 2 – 5 degrees. We imaged VSDI signals from two monkeys, and GCaMP signals from one monkey. We performed Epi-fluorescence imaging using the following filter sets: GCaMP, excitation 470/24 nm, dichroic 505 nm, emission 515 nm cutoff glass filter; VSDI, excitation 620/30 nm, dichroic 660 nm, emission 665 nm cutoff glass filter). Illumination was obtained with an LED light source (X-Cite120LED) for GCaMP or a QTH lamp (Zeiss) for VSDI imaging. Data acquisition was time locked to the animal’s heartbeat. The sampling frequency for GCaMP imaging is at 20 Hz, and for VSDI imagine it is at 100 Hz.

### Visual stimuli

The stimulus patterns were large-filed (8-degree radius) bandpassed noise patterns (centered at 3 cycles per degree), independently generated for each trial. The pattern was chosen because 1) it was used in previous studies that we compared our measurement results to; and 2) the patterns activated large V1 responses in humans. The patterns were presented with 12 different temporal conditions. The first six temporal conditions were a single pulse a stimulus presented with various durations (20, 40, 80, 160, 320, 640 ms). The last six temporal conditions were two stimulus pulses with varying inter-stimulus intervals. Each stimulus pulse lasts 160 ms, and the inter-stimulus intervals took values of 20, 40, 80, 160, 320, 640 ms.

### Data pre-processing

For each trial of the VSD experiment, we performed three pre-processing steps before extracting stimulus-evoked responses. First, we computed the averaged VSDI response dynamics to two blank trials (no stimulus). Then we subtracted this averaged time course from VSD dynamics measured in each stimulus-present trial. This step gets rid of noise that did not depend on stimulus, but was shared across all trials.

Next, for each trial, there was a time delay between stimulus onsets and response onsets. This is because visual signals take time to travel from animals’ eyes to their superficial layer of V1. We can either incorporate this delay into our models, or we can get rid of the delay by shifting response onsets to an earlier time point to match stimulus onset. Because this delay between response and stimulus onsets is not of interest to our analysis, we took the latter approach, and shifted the response time course forward by 60ms (for each trial). 60ms is determined by visualization, as well as on model cross-validation results that we will later describe in the Method section.

Third, for each time course, we subtracted the first entry from the entire time course. In other words, we set the beginning of each response time course to 0. As a consequence, the pre-processed VSDI dynamics are relatively similar at the beginning of the trials, and become more diverse towards the end of the trials. GCaMP data pre-processing takes the same steps as VSDI pre-processing.

### Two-component linear model

This model consists of a fast and a slow component, and each component is parameterized by a distinct set of basis functions. The fast component is designed to capture stimulus-evoked dynamics, and the fast component consists of a filter *f*(*t*) convolved with a stimulus time course *s*(*t*). Filter was parameterized as a weighted sum of basis functions, {*f*_*i*_(*t*)}. Each basis is a raised cosine function the input of which is a logarithmically warped time course [30]. The logarithmic warping intends to capture observations that neuronal dynamics are fast right after stimulus onsets, and gradually slow down over time [35]. We estimated the weights {*u*_*i*_} for each basis function. The slow component of the model *g*(*t*) intends to capture slow VSD dynamics. The slow component is parameterized using a different set of (slower) basis functions {*g*_*j*_(*t*)}, and we estimate weights {*v*_*i*_} for each of these bases. Differs from the fast component, the slow component does not depend on the stimulus, and is not convolved with the stimulus time course. Overall, the model can be summarized using a single equation:

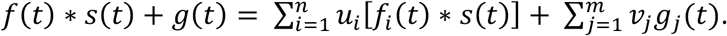

The set {*g*_*j*_(*t*)} is designed to be different from {*f*_*i*_(*t*)} in three ways to prevent trade-offs between the two components. First, the two sets of basis functions have different time scales, the fast basis functions cover a range up to 0.3 second for VSDI signals, and the slow basis functions cover a range of 1.2 seconds. Second, the fast basis functions are convolved with stimulus time courses, and the slow basis functions are directly added to the fast response without interacting with the stimulus. Third, the set {*f*_*i*_(*t*)} has fast-varying basis functions followed by slow ones, whereas in the slow-basis set {*g*_*j*_(*t*)} start slow but are followed by faster dynamical variations (see Figure S2B). This is because during the experiment, trial onsets are set to be time-locked to the animal’s heartbeat, and we re-set the initial point of each trial of VSDI dynamics to 0. Due to this initial alignment, VSDI time courses tend to vary more diversely toward the end of each trial, and this variation can be captured by denser and faster-varying basis functions in {*g*_*j*_(*t*)}.

Overall, this basis approach is simple -- the weight for each basis linearly contributes to the predicted overall VSDI responses and can be estimated via a closed-form least-square minimization. The approach is also flexible -- the set of basis functions for the fast component can flexibly approximate common shapes of neuronal filters, and the set of slow basis functions can flexibly approximate different slow variations in VSDI time courses.

### Two-component delayed normalization model

The two-component delayed normalization model has the same structure as the two-component linear model, and the slow component for this model is implemented the same way as the two-component linear model.

The fast component consists of a delayed normalization model [16]. The model has a numerator, and a denominator. The numerator consists of a stimulus time course *s*(*t*) convolved with a filter *f*_1_(*t*), which is parameterized using a set of basis functions, as the linear model. The denominator of the model consists of two parts, a non-negative constant *σ* that prevents the denominator from being 0 (therefore the predicted outcome is un-defined), and another filter *f*_1_(*t*) that convolves with the same stimulus time course *s*(*t*). The filter *f*_1_(*t*) is parameterized as a difference between two Gamma functions (see [16]), this parameterization makes additional assumption on the shape of the denominator filter, and better constraints parameter estimations. Overall, the delayed normalization model can be written as

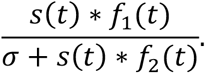

The estimated denominator filter is typically slower than the numerator filter in practice, which gives rise to the prediction of slower dynamics at low stimulus contrast or at brief stimulus durations. This is because of the division – the suppressive effect of the denominator is small when stimulus drive (numerator) is also small, and the model dynamics are dominated by the numerator response. If stimulus input becomes large (e.g. high image contrast), suppressive effect from the denominator exponentially grows, and the denominator starts to dominate and speed up the dynamics (e.g. response decays sharply after the initial transient).

Using trial-averaged data, we estimated two-component delayed normalization parameters. We separated the parameters into two batches, parameters for delayed normalization, and weights for the additional slow basis functions (for the slow data component). We alternated between the two sets of parameters, and used coordinate descend to find model parameters.

### Alternative models to extract stimulus-evoked components in VSD and GCaMP

#### Linear boxcar with one filter per stimulus

For a stimulus condition, this model assumes the measurement time course can be separated into two additive components, one relatively fast and one slow. The fast component is assumed consisting of a convolution between a scaled linear filter (parameterized as a gamma function) and a stimulus time course, and the slow component is assumed linear, and is modeled as a scalar multiplies the time course:

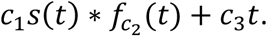

We fit three parameters for this model, a scaler *c*_1_ for the fast component, a parameter *c*_1_ that governs the shape of the fast filter, and a scaler *c*_*_ for the slow component. Because these parameters nonlinearly interact with each other, we used matlab built-in function *fminsearch*.*m* to find model parameters.

#### Exponential boxcar with one filter per stimulus

This model shares a similar structure with the previous model, except that the slow dynamics is modeled as an exponential (typically a decay), instead of a linear function. So this model can be expressed as:

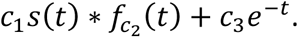

Notice that both in this model and in the previous model, we fitted a different filter for each stimulus condition. The other two models in Supplementary Figure X are the same as these two models, but we assume a single fast filter is shared across stimulus conditions for each model.

### Model comparison

We performed leave-one-condition out cross-validation to compare between two-component linear and two-component delayed normalization model. For this cross-validation, we fit the entire model (either two-component linear or normalization model) to the trial-averaged left-in condition (11 out of 12 temporal conditions), and used the fitted model parameters that pertained to the fast components to make predictions to the fast component of the left-out condition. We combined the predicted fast component to the left-out condition with an additionally fitted slow-component to compute the overall variance explained.

### Comparing between VSD, GCaMP and fMRI dynamics

In Figure 3, we compared stimulus-evoked component in VSD and in GCaMP data, to temporal dynamics measured using fMRI [15] and ECoG [16]. The fMRI experiment shares the same stimulus design and a comparable set of temporal conditions, so we emphasized the comparison between fMRI and VSD data. The ECoG data are averaged across different stimulus patterns, therefore we only used it to demonstrate that its temporal properties are consistent with that measured in fMRI.

For fMRI data, one number is extracted for each stimulus condition, this is because fMRI hemodynamics are many times slower than neuronal dynamics, and estimating the time course of neuronal dynamics would be extremely challenging. The single number extracted for each condition can be viewed as the sum of neuronal responses for that stimulus condition.

For ECoG, delayed normalization model was fit to ECoG broadband responses measured in multiple humans’ V1. Broadband responses were thought to relate to LFP, and were correlated with fMRI responses. To make ECoG predictions comparable to the fMRI measurements, each predicted ECoG time course was summed for a stimulus condition, and an overall scaling factor was fit to the summed responses to make ECoG and fMRI response span the same range.

To make stimulus-evoked VSD and GCaMP component comparable to both fMRI and ECoG broadband responses, a two-component normalization model was fit to the trial-averaged measurement time courses. For each condition, we subtracted the estimated slow component away from the data, and summed remaining time courses to get 12 numbers. We compared the 12 numbers obtained from VSD and GCaMP to the prediction of a linear model, as well as to fMRI and ECoG responses.

## Supplementary Figures

**Figure S1.**
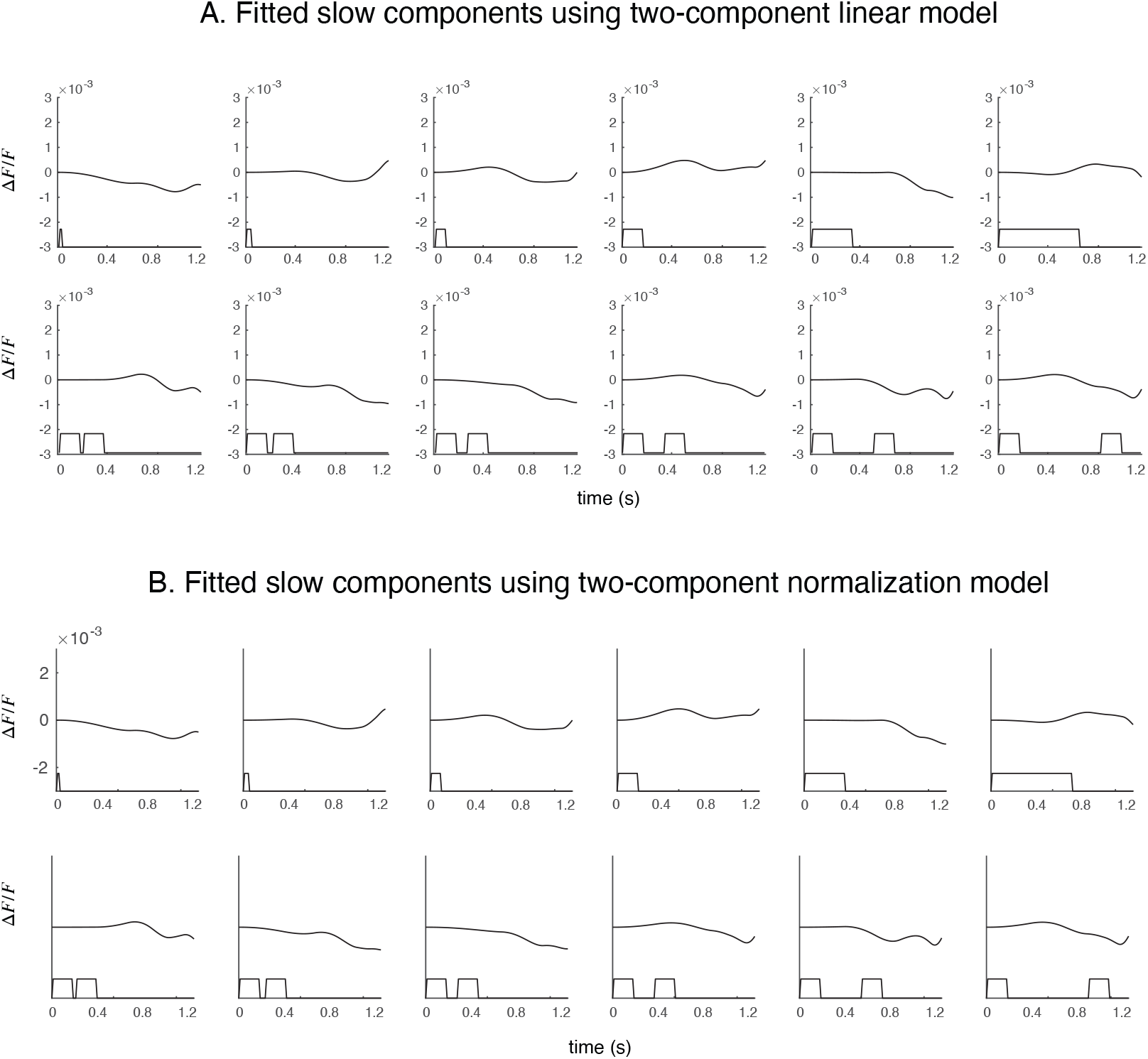
Slow variations in VSD do not seem to depend on stimulus. We visualize slow dynamics fit to VSD time courses using the two-component linear model (panel A), and using the two-component delayed normalization model (panel B). The slow dynamics are fit with a set of bases functions, and we estimate a weight for each basis (see Figure S1). The estimated slow dynamics using both models don’t seem to systematically vary with different stimulus conditions.

**Figure S2.**
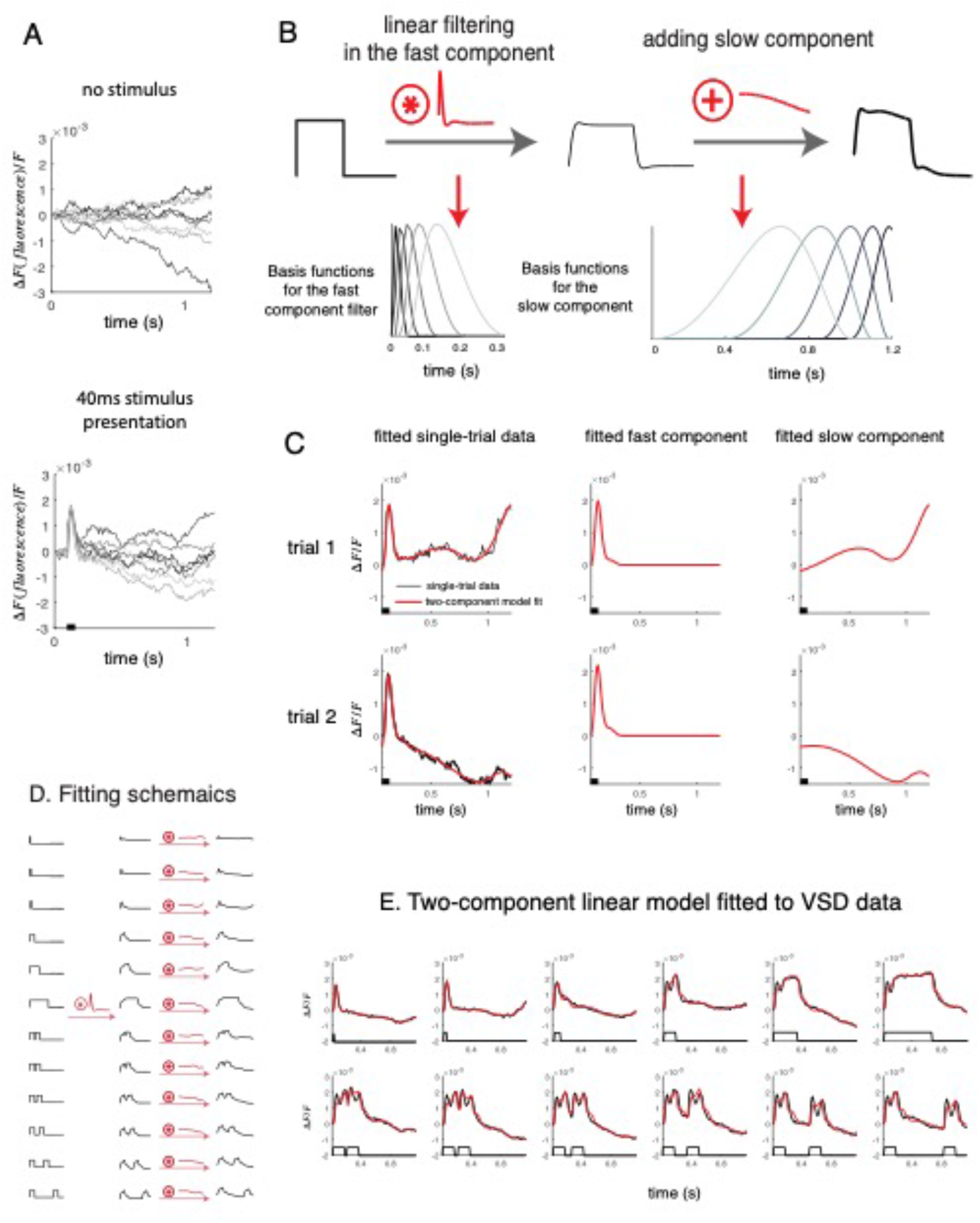
Two-component linear model: schematics. Two-component linear model. A. Different trials of VSD time courses responding to no stimulus (upper panel), and to a brief stimulus pulse (lower panel). Across different trials, there seems to be a dynamical component that closely tracks the stimulus time course, and is consistent across trials. Additionally, there are other slower dynamics that vary from trial to trial, even at the presentation of the same stimulus time course. B. The two-component linear model consists of a fast component (a filter convolves with stimulus time course), and a slow component that are additively combined with the fast time course. Both the fast filter and the slow component are parameterized using (different) sets of basis functions. C. Fitting the two-component linear model to two example trials. We can see that between the two trials, the fitted fast components are similar between trials, and the fitted slow component vary across trials. D. For our analysis in Results in the main text, we fitted a single set of fast filters, and distinct set of slow dynamics (one for each temporal condition) to trail-averaged VSD dynamics. E. The fitted dynamics closely match with the trial-averaged data for each stimulus condition.

**Figure S3.**
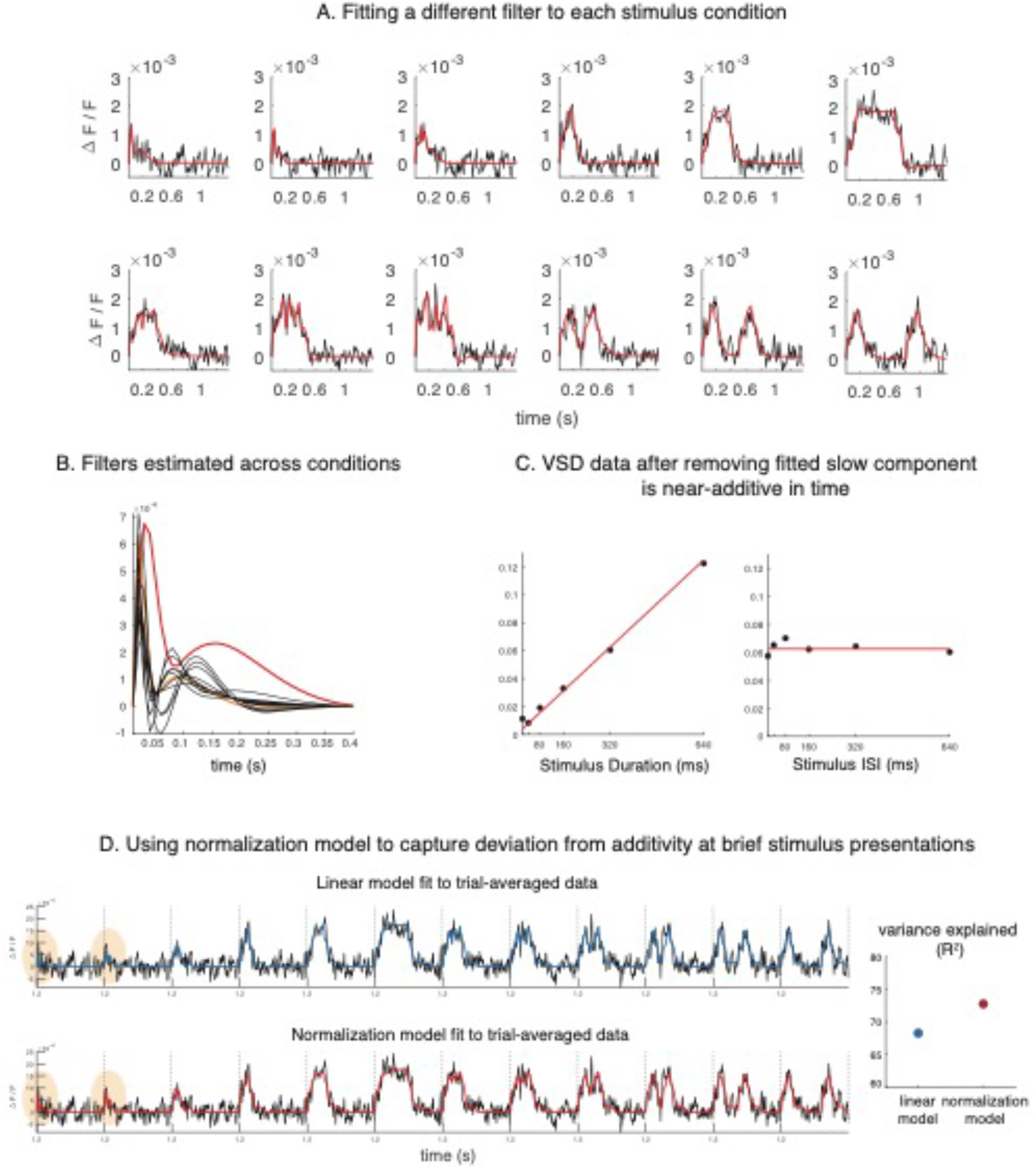
An additional set of VSD data. Analyzing an additional set of VSD data leads to conclusions similar to the main text. A. We extracted stimulus-evoked signals using a two-component linear model. A different filter was fit to each stimulus condition. B. Filters for very brief stimulus presentation durations (red) seem to have a different shape, compared to the filters estimated for the rest of the conditions. C. Stimulus-evoked VSD dynamics are near-additive. D. Delayed normalization model leads to improved description of stimulus-evoked VSD dynamics, compared to the linear model.

**Figure S4.**
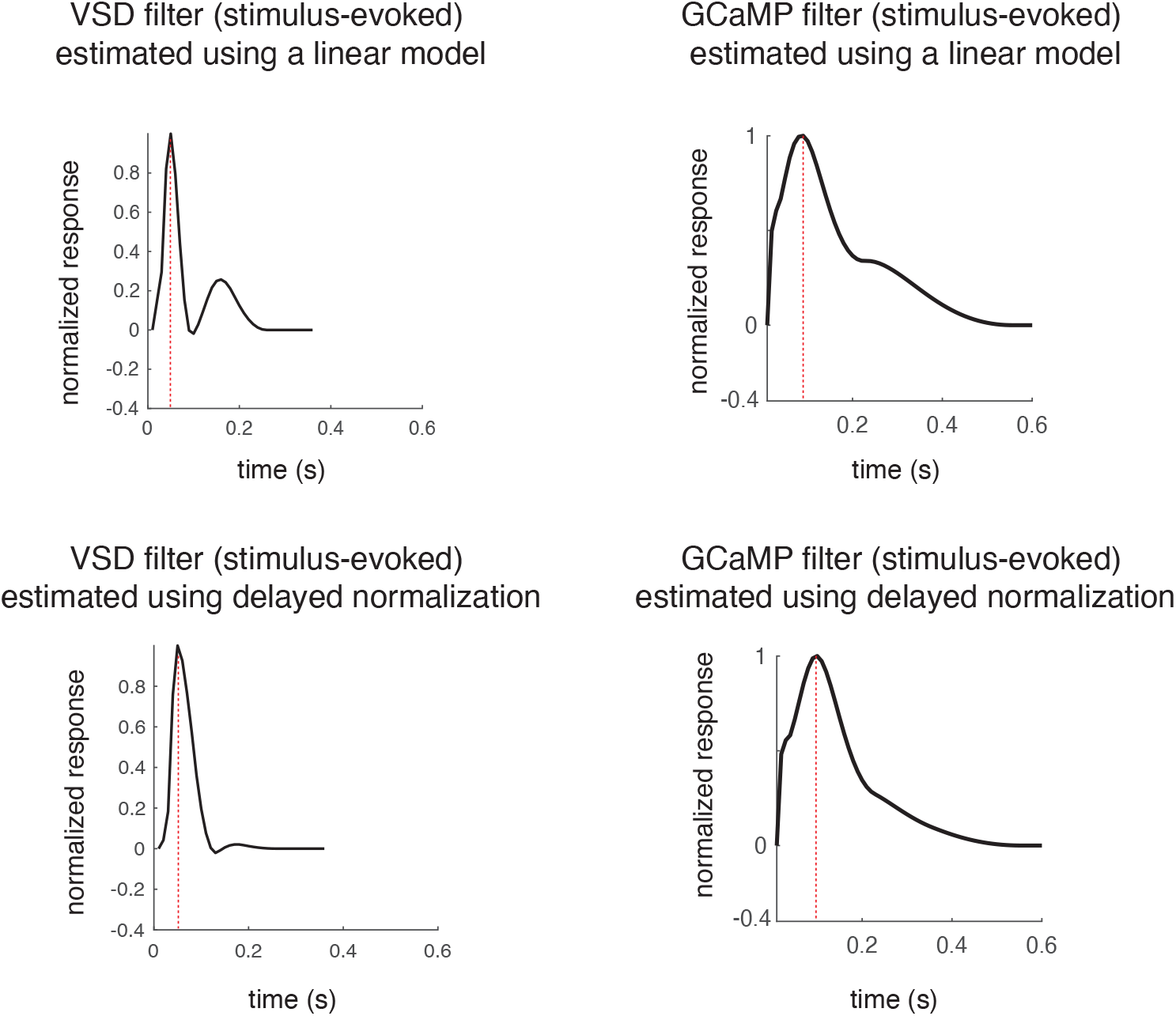
Estimated filters for stimulus-evoked VSD and GCaMP dynamics. For both VSD and GCaMP, filters estimated using two-component delayed normalization seem simpler in shape. This is because part of the filter dynamics (estimated using two-component linear model) are absorbed into the normalization dynamics. VSD filters have shorter time to peak (50 ms), compared to GCaMP filters (∼100 ms). The time to peak in each filter seems to be invariant to which model was used.

**Figure S5.**
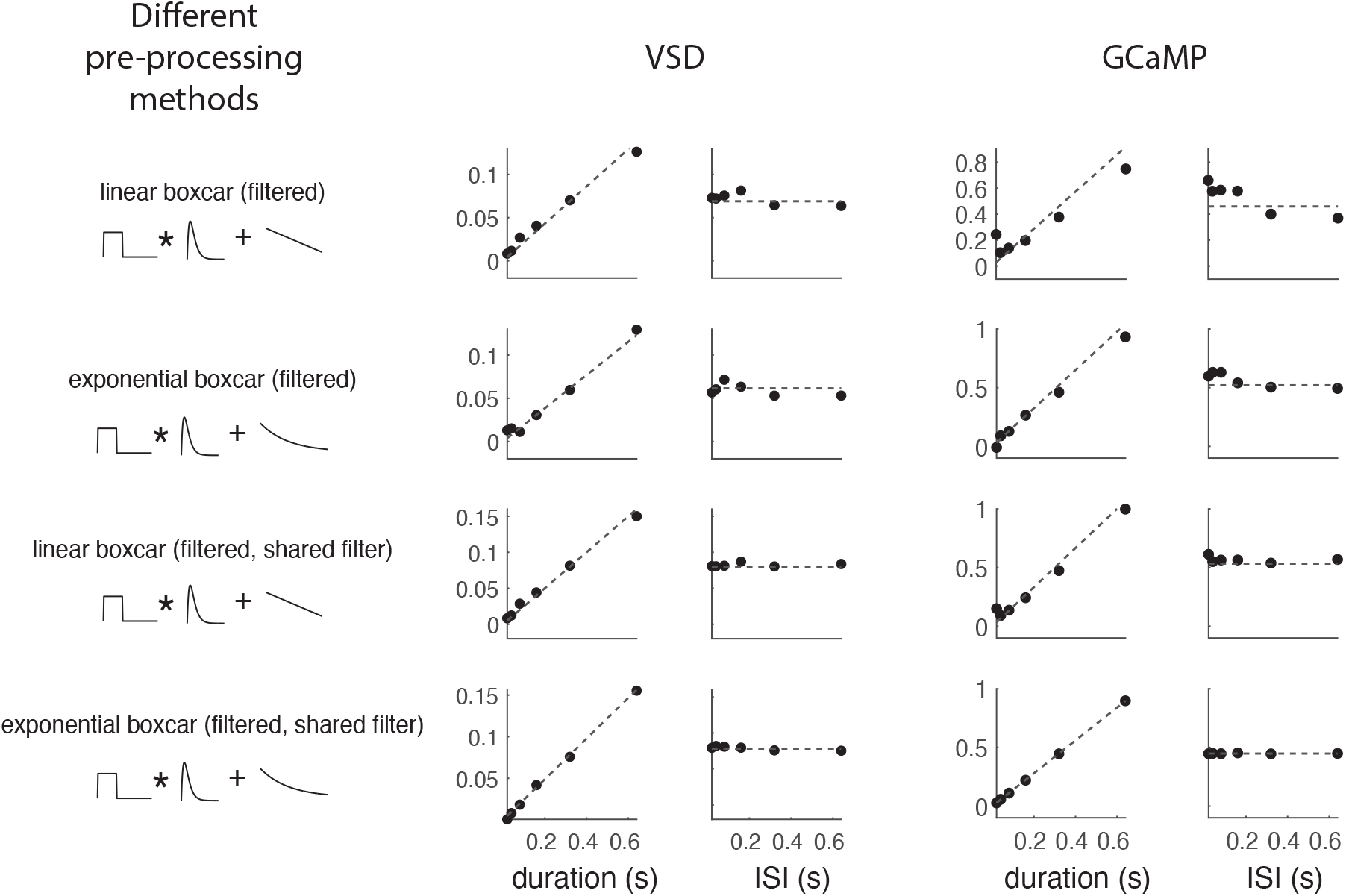
Confirm near-additive temporal dynamics using other methods to extract stimulus-evoked signals. To examine whether near-additive VSD and GCaMP dynamics is a result of our data extraction methods, we used 4 other different methods to extract stimulus-evoked components from both types of signals, and reached similar conclusions. The first model is a linear boxcar model, and we assume signal dynamics can be approximated by a sum of fast component (a stimulus time course convolved with a filter), and a linear trend. In the second model, we assume the same fast component, but the slow component is approximated with an exponential decay. Notice that in the first two models, a distinct filter was estimated for each stimulus condition. The last two models are the same as previous two models, except that a single filter was estimated for all stimulus conditions.

